# Central amygdala mineralocorticoid receptors modulate alcohol self-administration

**DOI:** 10.1101/2020.05.19.105262

**Authors:** Viren H. Makhijani, Preethi Irukulapati, Kalynn Van Voorhies, Brayden Fortino, Joyce Besheer

## Abstract

The mineralocorticoid receptor (MR) is an emerging target in the field of alcohol research. The MR is a steroid receptor in the same family as the glucocorticoid receptor, with which it shares the ligand corticosterone in addition to the MR selective ligand aldosterone. Recent studies have shown correlations between central amygdala (CeA) MR expression and alcohol drinking in rats and macaques, as well as correlations between aldosterone and alcohol craving in individuals with alcohol use disorder (AUD). Additionally, our previous work demonstrated that systemic treatment with the MR antagonist spironolactone reduced alcohol self-administration and response persistence in both male and female rats. This study examined if reductions in self-administration following MR antagonist treatment were related to dysregulation of MR-mediated corticosterone negative feedback. Female rats treated with spironolactone (50 mg/kg; IP) showed increased plasma corticosterone following self-administration which correlated with reduced alcohol self-administration. Next, local microinjection of the MR-selective antagonist eplerenone was used to identify the brain-regional locus of MR action on alcohol self-administration. Eplerenone infusion produced dose-dependent reductions in alcohol self-administration in the CeA, but had no effect in the dorsal hippocampus. Finally, to assay the functional role of CeA MR expression in alcohol self-administration, CeA MR was knocked down by antisense oligonucleotide (ASO) infusion prior to alcohol self-administration. Rats showed a transient reduction in alcohol self-administration 1 day after ASO infusion. Together these studies demonstrate a functional role of CeA MR in modulating alcohol self-administration and make a case for studying MR antagonists as a novel treatment for AUD.

## 1. INTRODUCTION

Alcohol use disorder (AUD) poses a significant public health concern, affecting approximately 14.4 million adults in the United States, with an estimated societal cost of $249 billion (Sacks et al., 2015, SAMHSA, 2018). However, despite widespread prevalence and significant societal cost, prescription data suggests about 9% of AUD patients receive pharmacological treatment, despite its proven efficacy (Mark et al., 2009, Kranzler and Soyka, 2018, Ray et al., 2019). Barriers to pharmacological treatment of AUD include factors such as lack of perceived efficacy by both patients and physicians, belief that AUD is managed by addiction specialists and not primary care physicians, and alcohol-related stigma (Finlay et al., 2017, Williams et al., 2018). As such there is a need for the development of novel AUD therapeutics that not are not only efficacious, but may spur widespread adoption.

One emerging pharmacological target is the mineralocorticoid receptor (MR), a steroid receptor closely related to the glucocorticoid receptor (GR), another pharmacological target for AUD (Vendruscolo et al., 2015, Shen, 2018). It has long been known that GRs and corticosterone, as part of the hypothalamic-pituitary-adrenal (HPA) axis, influence alcohol self-administration and mediate alcohol dependence (Fahlke et al., 1994a, Fahlke et al., 1994b, Fahlke et al., 1995, Koenig and Olive, 2004, Vendruscolo et al., 2012). Recent work from a multi-species study has provided evidence for a role of MR and its’ ligand aldosterone in alcohol consumption (Aoun et al., 2018). Male rhesus macaques and Wistar rats with alcohol experience showed inverse correlations between central amygdala (CeA) MR expression and measures of alcohol drinking, while human AUD patients showed correlations between circulating aldosterone and alcohol craving (Aoun et al., 2018). Our previous findings built upon this study and demonstrated that the MR antagonist spironolactone, an FDA approved drug indicated for managing hypertension, reduced alcohol self-administration and response persistence in male and female rats (Makhijani et al., 2018). Together, these studies have begun to outline a relationship between MR and alcohol drinking; however, the functional role of CeA MR in alcohol drinking is not fully understood and it is unknown if altering CeA MR expression would alter alcohol drinking.

The goal of the present study was to build on this previous work by first assessing if MR antagonism may mediate reductions in alcohol self-administration by impairment of MR-mediated corticosterone negative feedback (Atkinson et al., 2008). Next, the second-generation MR antagonist eplerenone, which has higher MR selectivity than spironolactone, was infused into the dorsal hippocampus (dHC), or CeA to identify the brain-regional locus of MR action on alcohol self-administration. These target regions were chosen as the dHC has the highest levels of MR expression in the brain (McEwen et al., 1968), and the CeA is where MR expression correlates with alcohol drinking (Aoun et al., 2018). Finally, to assay the functional role of CeA MR expression in modulating alcohol self-administration, CeA MR was knocked down by antisense oligonucleotide (ASO) infusion prior to alcohol self-administration.

## 2. METHODS AND MATERIALS

### 2.1. Animals

72 Long-Evans rats (60 female and 12 male; Envigo-Harlan, Indianapolis, IN) arrived at 7 weeks old and were housed under a 12-hour light/dark cycle (7:00 am/pm). Female rats were used for all self-administration studies. Our previous studies reported similar effects of MR antagonism on alcohol self-administration between sexes (Makhijani et al., 2018). In Experiments 1 and 2 rats were single-housed throughout the study, in Experiments 3 and 4 rats were double-housed until surgery and then single-housed through the rest of the study. Prior to all experiments rats were handled for 1-2 minutes for 5 days. All experiments were conducted during the light cycle. Animals were under the care of the UNC-Chapel Hill Division of Comparative Medicine veterinary staff. All procedures were carried out in accordance with the NIH Guide for Care and Use of Laboratory Animals, and institutional guidelines. All protocols were approved by the UNC Institutional Animal Care and Use Committee (IACUC). UNC-Chapel Hill is accredited by the Association for Assessment and Accreditation of Laboratory Animal Care (AAALAC).

### 2.2. Behavioral and surgical protocols

#### 2.2.1. Self-administration apparatus

Self-administration was conducted in operant chambers (Med Associates, Georgia, VT) within sound-attenuating cabinets which contained an exhaust fan to provide ventilation and mask external noise. Chambers were equipped with two retractable levers on opposite sides of the chamber (left and right), and a cue light was located above each lever. When the response requirement was met on the left (active) lever, a cue light (directly above the lever) and a stimulus tone were presented for the duration of the alcohol reinforcer delivery (0.1 mL of solution into a well on the left side of the chamber across 1.66 s via a syringe pump). Responding during reinforcer delivery and on the right (inactive) lever was recorded, but had no programmed consequences. Chambers were also equipped with 4 parallel infrared beams across the bar floor to measure general locomotor activity during the session. The number of beam breaks for the entire session was totaled and divided by the session length (30 min) to calculate the locomotor rate (beam breaks/min).

#### 2.2.2. Alcohol self-administration

Rats were trained to self-administer a 20% (v/v) alcohol solution (20A) on a fixed ratio 2 (FR2) reinforcement schedule across 30-minute sessions, five days a week (M-F) via sucrose fading as described in (Makhijani et al., 2018, Randall et al., 2017). Sucrose fading began with self-administration of 2% alcohol/10% (w/v) sucrose (2A/10S), then 5A/10S, 10A/10S, 10A/5S, 15A/5S, 15A/2S, and 20A/2S on subsequent sessions, ending with 20A which remained the reinforcer through the remainder of the study. Rats in Experiments 1 and 2 had approximately 8 weeks of self-administration training and were used in an unrelated study (i.e., involved exposure to a single stressor and self-administration was unaltered (unpublished)) two months prior to the initiation of this study.

#### 2.2.3. Surgical procedures and microinjections

For Experiment 2, rats were anesthetized with isoflurane (3-5% in 98% oxygen; Baxter, Deerfield, IL) and received implantation of 22-gauge guide cannulae (P1 Technologies, Roanoke, VA) aimed to terminate 2 mm above the central nucleus of the amygdala (n=18; CeA; bilateral coordinates: AP −2.5, ML ±4.2 mm, DV −5.8 mm) or the dorsal hippocampus (n=14; bilateral coordinates: AP −2.5, ML ±1.5 mm, DV −1.4 mm). For Experiments 3 and 4, anesthetized rats received implantation of 26-gauge guide cannulae (P1 Technologies) aimed to terminate 2 mm above the CeA (Experiment 3 n=12 males, Experiment 4 n = 24 females). Coordinates were based on (Paxinos and Watson, 2013).

Site-specific bilateral microinjections were delivered by a microinfusion pump (Harvard Apparatus, Holliston, MA) using 5.0 μl Hamilton syringes connected to 28 (Experiment 2) or 33-gauge (Experiments 3 & 4) injectors that extended 2 mm below the guide cannulae (P1 Technologies). Rate of infusions were 0.5 μl/side across 1 minute (Experiment 2-eplerenone) or 5 minutes (Experiments 3 & 4 - ASO). The injector(s) remained in place for an additional 2 minutes (eplerenone) or 5 minutes (ASO) after the infusion to allow for diffusion. Additional microinjection procedures are described in detail in (Besheer et al., 2012, Jaramillo et al., 2018).

At the end of Experiments 2, 3, and 4 brain tissue was stained with cresyl violet to verify cannulae placement. Only data from rats with both cannulae/injector tracts determined to be in the target brain regions were used in the analyses.

### 2.3. Tissue collection and molecular analyses

#### 2.3.1. Blood collection and corticosterone EIA

Tail blood was collected immediately after alcohol self-administration on the spironolactone test day in Experiment 1 for assessment of plasma corticosterone. Blood was collected into heparinized tubes and immediately centrifuged at 4°C for 5 minutes at 2000 rcf. Plasma supernatant was then collected and stored at −80°C until analysis. 5 μL plasma samples were then analyzed in duplicate using a commercially available colorimetric EIA kit (ArborAssays, Ann Arbor, MI) according to the manufacturer’s instructions.

#### 2.3.2. Tissue collection and preparation

Rats were sacrificed by rapid decapitation under deep isoflurane anesthesia (Baxter, Deerfield, IL). Brains were flash-frozen in isopentane chilled on dry ice and stored at - 80°C until tissue punching. Frozen brains were sectioned coronally by cryostat and bilateral 1.2 mm diameter punches were taken of central amygdala (CeA; AP −1.8 to - 2.8). One tissue punch was processed for western blotting and one was processed for qPCR as follows. Western blot samples were homogenized by sonication (Branson SLPe; Emerson Industrial, St. Louis, MO) in homogenization buffer (10mM Tris-HCl, 1% (w/v) SDS, 1% (v/v) HALT protease inhibitor, 5mM EDTA). Protein concentration from lysates were determined using a BCA assay (Thermo-Fisher; Rockford, IL). RNA was extracted from qPCR samples using the RNeasy Mini Kit (Qiagen, Venlo, Netherlands) according to manufacturer’s instructions and eluted in 30uL of nuclease free water (Life Technologies Corp., TX, USA). RNA concentration and purity for each sample were determined using a spectrophometer (Nanodrop 2000, Thermo-Fisher).

#### 2.3.3. Western blotting

10 μg of protein lysate was mixed with LDS sample buffer (Life Technologies, Carlsbad, CA), reducing agent (Life Technologies) and sterile water and denatured at 70°C for 5 minutes. Protein samples were then separated using an 18-well 4-20% Criterion TGX gel (Bio-Rad, Hercules, CA) in a Bio-Rad western blot apparatus. All samples were run on one gel. After electrophoresis, proteins were transferred to PVDF membrane using the iBlot system (Thermo-Fisher). Blots were blocked in 3% non-fat dry milk (Nestle, Solon, OH) in 0.1% PBST (0.1% Tween-20 in PBS) prior to incubation with antibodies against the mineralocorticoid receptor (1:400, Lot #’s: 3237523, 3083584; MABS496, EMD Millipore, Temecula, CA) and the loading control actin (1:5000, Lot#: 3086655; MAB1501, EMD Millipore). After incubation with primary antibody, blots were washed and incubated with an HRP anti-mouse secondary antibody (1:10,000, Lot#: X0328; Vector Labs, Burlingame, CA). Blots were then developed in chemiluminescent substrate (ECL-Prime, GE Healthcare, Little Chalfont, UK) and imaged using an Imagequant LAS 4000 (GE Healthcare). Band optical density measurements were collected and analyzed using ImageQuantTL software (GE Healthcare).

#### 2.3.4. Reverse transcription and qPCR

RNA was reverse transcribed into circular DNA (cDNA) using the QuantiNova Reverse Transcription Kit (Qiagen, Venlo, Netherlands) according to manufacturer’s instructions. cDNA was diluted 1:10 with water and stored at −20°C before qPCR experiments. All qPCR reactions were run with a QuantStudio3 real-time PCR machine (ThermoFisher) was used for all experiments. Using a 96-well plate, each sample was run in triplicate using 10uL total volume per well with the following components: PowerUp Syber green dye (ThermoFisher), forward and reverse primers (Eton Biosciences Inc., NC, USA), and cDNA template. The PCR was run with an initial activation for 10 min at 95°C, followed by 40 cycles of the following: denaturation (95°C for 15s), annealing (60°C for 30s), and extension (72°C for 45s). Melt curves were obtained for all experiments to verify synthesis of a single amplicon. Primers used were mineralocorticoid receptor (NR3C2): F: 5’-GAT CCA GGT CGT GAA GTG GG-3’, R: 5’-AGA GGA GTT GGC TGT TCG TG-3’; β-actin (ACTB) (loading control): F: 5’-CTA CAA TGA GCT GCG TGT GGC-3’, R: 5’-CAG GTC CAG ACG CAG GAT GGC-3’.

#### 2.3.5. Oligonucleotides, drugs, and reagents

Oligonucleotide sequences were sourced from previous literature (Sakai et al., 2000, Johnson and Greenwood-Van Meerveld, 2015) which had functionally validated MR knockdown using these sequences. 18-mer phosphonothioate oligonucleotide sequences were: MR antisense (ASO): 5′-TTC CAT GTC TAG GCC TTC-3′, MR scrambled (SCR): 5′-CAT TTT GAA GGT TCC GGT-3′. Oligonucleotides were synthesized by Eurofins Genomics (Louisville, KY) and supplied as dried, salt-free stocks that were suspended in artificial cerebrospinal fluid (ACSF; Tocris, Bristol, UK) for microinjection.

Spironolactone (Lot #’s: MKCD7812, MKCG6303; Sigma-Aldrich, St. Louis, MO) and eplerenone (Lot #’s: 1B/209648, 1B/205653, 1B/210844; Sigma-Aldrich, St. Louis, MO) were suspended in 45% 2-hydroxypropyl-β-cyclodextrin (Sigma-Aldrich) for injection and microinjection. Alcohol (95% (v/v), Pharmaco-AAPER, Shelbyville, KY) and sucrose (Great Value, Bentonville, AR) were diluted with tap water for all self-administration sessions.

### 2.4. Experimental procedures

#### 2.4.1. Experiment 1: Role of inhibited glucocorticoid negative feedback in spironolactone suppression of alcohol self-administration

The purpose of this experiment was 1) to quantify the effect of MR antagonism by spironolactone on glucocorticoid negative feedback, as mineralocorticoid receptor signaling is important in maintaining basal corticosterone levels (Joels and de Kloet, 2017), and 2) to determine if changes in corticosterone may underlie spironolactone-induced decreases in alcohol self-administration.

Following establishment of stable alcohol self-administration (approximately 40 sessions), rats (n = 18/group) received spironolactone (0, 50 mg/kg; IP; 1 mL/kg) 30 minutes prior to an alcohol self-administration session. We have previously demonstrated that this dose of spironolactone reduces alcohol self-administration in male and female rats (Makhijani et al., 2018). Immediately after the 30-min self-administration, blood was collected for analysis of plasma corticosterone as described above. Testing was conducted during a single self-administration session using a between-subjects design.

#### 2.4.2. Experiment 2: Effect of intra-dorsal hippocampus and intra-central amygdala eplerenone on alcohol self-administration

The purpose of this study was to determine the brain regional locus of MR antagonism-induced reduction of alcohol self-administration by using site specific microinjection of the selective MR antagonist eplerenone.

Following Experiment 1, rats were implanted with bilateral cannulae aimed at the dHC (n = 14) or CeA (n = 18). To determine the role of CeA and dHC MR in alcohol self-administration, rats received a bilateral infusion of eplerenone (0, 100, 1000, 5000 ng/0.5 μL/side) immediately before an alcohol self-administration session. A within-subject design was used such that each rat received each treatment in a random order and doses were equally represented on each test day. Test days were on Tuesdays and Thursdays with standard self-administration sessions on Monday, Wednesday, and Friday. Alcohol lever responses had to be at least 80% of baseline (average responding of the 2 sessions preceding the initiation of testing) self-administration on the days preceding a test day in order for a rat to be tested - all rats met this criterion.

#### 2.4.3. Experiment 3: Confirmation of gene knockdown using ASO infusion

The purpose of this experiment was to determine the longevity of gene knockdown from infusion of a validated MR ASO construct (Sakai et al., 2000, Johnson and Greenwood-Van Meerveld, 2015).

Naïve rats were implanted with bilateral cannulae aimed at the CeA and allowed 1 week for recovery. Rats (n = 3/group) were then infused with either ASO or SCR and sacrificed by rapid decapitation under deep isoflurane (Baxter) anesthesia either 2 or 7 days later. Brains were collected for confirmation of gene and protein knockdown by qPCR and western blot, respectively.

#### 2.4.4. Experiment 4: Effect of central amygdala mineralocorticoid receptor knockdown on alcohol self-administration

The purpose of this experiment was to assess the functional role of CeA mineralocorticoid receptor tone in alcohol self-administration. Following establishment of stable alcohol self-administration (15 sessions at maintenance concentration), rats (n = 24) were implanted with bilateral cannulae aimed at the CeA and allowed 1 week for recovery and then returned to alcohol self-administration. After 5 alcohol self-administration sessions, rats received a bilateral infusion of ASO or SCR (n = 12/group) 3 μg/0.5 μL/side. Self-administration was withheld on this infusion day. 24 hours after the infusion, alcohol self-administration continued for one week (5 sessions).

### 2.5. Data analysis

#### 2.5.1. Alcohol self-administration

In Experiment 1, alcohol lever responses, inactive lever responses, alcohol intake (g/kg, estimated from bodyweight and number of reinforcers received), and locomotor rate were compared by student’s t-test. Plasma corticosterone following alcohol self-administration was compared by student’s t-test and relationships between plasma corticosterone and alcohol self-administration were evaluated by Spearman’s rank order correlation. In Experiment 2, alcohol lever responses, inactive lever responses, alcohol intake, and locomotor rate were compared by one-way RM-ANOVA with eplerenone dose as the within-subjects repeated measure. Cumulative alcohol lever responses are examined by two-way RM-ANOVA with eplerenone dose and time as within-subjects repeated measures. In Experiment 4, rats were grouped by baseline self-administration measures taken from the 5 post-cannulation self-administration session prior to oligonucleotide infusion. Alcohol lever responses, inactive lever responses, and locomotor rate were compared by two-way RM-ANOVA with oligonucleotide as the between-subjects measure, and session as the within-subjects repeated measure. Alcohol consumption is represented as an average across all post-infusion self-administration sessions and compared by two-way RM-ANOVA with oligonucleotide as the between-subjects measure, and pre/post-infusion as the within-subjects measure. For all experiments using a between-subjects design, groups were counterbalanced by alcohol self-administration history (average session alcohol lever responses and alcohol intake) across the past week (5 sessions) of self-administration.

#### 2.5.2. Western blotting

Mineralocorticoid receptor band densities were normalized to actin band density to account for loading variation, normalized protein levels in the ASO group were then expressed as a percent of the average normalized protein level for the pooled SCR groups (i.e. 2 day and 7 day as student’s t-test showed no difference in MR expression normalized to actin between timepoints). Due to unequal standard deviations between groups as detected by Brown-Forsythe test, protein expression between SCR and ASO groups were compared by Welch’s ANOVA and Dunnett’s T3 multiple comparisons test.

#### 2.5.3. qPCR

The threshold cycle (CT) of each target product was determined by software and the ΔΔCT method was used to calculate the percent change relative to control (CON). The ΔΔCT of NR3C2 (mineralocorticoid receptor) was then normalized to the ΔΔCT of ACTB (beta actin) and all values were expressed as a percentage of the pooled (2 and 7 days) SCR group’s normalized NR3C2 levels. mRNA levels were compared by one-way ANOVA. Given the unequal standard deviation between the SCR and 2-day ASO groups (confirmed by F-Test) and low sample size, a follow up Welch’s t-test was conducted to compare mRNA levels between the SCR and 2-day ASO groups and confirm findings from the western blot.

## 3. RESULTS

### 3.1. Experiment 1: Role of inhibited glucocorticoid negative feedback in spironolactone suppression of alcohol self-administration

The purpose of this experiment was to quantify the effect of MR antagonism on plasma corticosterone levels following alcohol self-administration and determine if increases in plasma corticosterone were related to reductions in alcohol self-administration. Blood was not collected from one animal in the spironolactone group and this animal was excluded from the analyses in Figs 1C&D.

**Figure 1.**
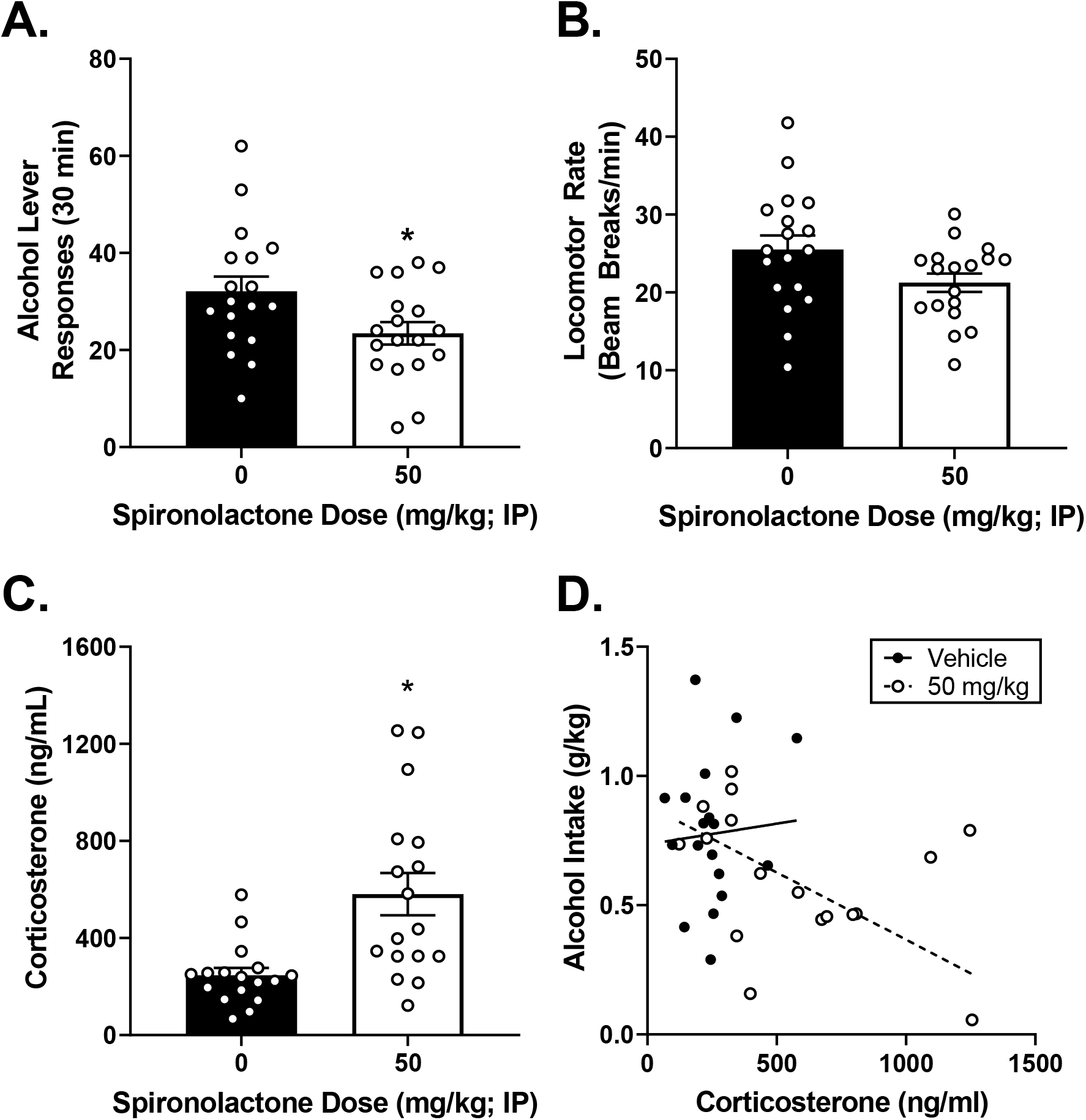
Spironolactone reduces alcohol self-administration and increases plasma corticosterone in female rats. (A) Spironolactone (50 mg/kg) reduces alcohol self-administration in female rats (n = 17/group). (B) Spironolactone has no effect on locomotor rate during the self-administration session. (C) Spironolactone treated animals have significantly higher plasma corticosterone following self-administration. (D) Plasma corticosterone correlates with reduced alcohol self-administration in spironolactone treated animals (R^2^ = 0.265, p = 0.037). *p < 0.05 versus vehicle.

#### 3.1.1. The MR antagonist spironolactone reduces alcohol self-administration in female rats, and increases plasma corticosterone following self-administration

Following treatment with the MR antagonist spironolactone, female rats showed significant reductions in alcohol self-administration (Fig 1A; t(34) = 2.28, p = 0.029), and alcohol intake (Table 1; t(34) = 2.08, p = 0.045). There were no effects on locomotor rate (Fig 1B) or inactive lever responses (Table 1). Plasma corticosterone was significantly elevated in the spironolactone treated animals (Fig 1C; t(33) = 3.72, p < 0.001).

**Table 1.** Inactive lever responses and alcohol intake for Experiment 1

| Group | Inactive Lever Responses | Alcohol Intake (g/kg) |
| --- | --- | --- |
| 0 mg/kg Spironolactone | 1.8 ± 0.4 | 0.79 ± 0.06 |
| 50 mg/kg Spironolactone | 1.6 ± 0.6 | 0.60 ± 0.06* |

\* - $p < 0.05$ versus 0 ng Eplerenone

#### 3.1.2. Elevated plasma corticosterone in spironolactone treated animals correlates with reduced alcohol self-administration

In order to determine if reductions in self-administration were related to increased plasma corticosterone, the relationship between the two measures was examined by Spearman’s rank order correlation given the non-linear nature of the relationship between corticosterone and behavior (Calabrese, 2008)(Fig 1D). There was a significant negative correlation between alcohol intake and plasma corticosterone in the spironolactone treated group (R^2^ = 0.265, p = 0.037), but not the control group (r = - 0.090, p = 0.723). This data pattern indicates that the effects of spironolactone on alcohol self-administration may be mediated by stimulation of the HPA axis through inhibition of glucocorticoid negative feedback.

### 3.2. Experiment 2: Effect of intra-dHC and intra-CeA eplerenone on alcohol self-administration

In order to determine the functional role of MR in alcohol self-administration, the MR antagonist eplerenone was infused into the dHC and CeA. 2 rats in the dHC group, and 6 rats in the CeA group were excluded from the study due to lost cannulae implants during the experiment.

#### 3.2.1. Intra-CeA but not intra-dHC eplerenone reduces alcohol self-administration

dHC cannulae verifications are represented in Fig 2A, 1 rat had cannula(e) outside the target region (depicted as open triangles) and was considered a miss and excluded from these analyses. Intra-dHC eplerenone had no effect on total alcohol lever responses, cumulative alcohol lever responses, inactive lever responses, locomotor rate (Fig 2B-E, respectively), or alcohol intake (Table 2). CeA cannulae verifications are shown in Fig 3A, 1 rat had cannula(e) outside the target region and was excluded from these analyses. In contrast, intra-CeA eplerenone significantly reduced alcohol self-administration as confirmed by a significant main effect of drug on total alcohol lever responses (Fig 3B; F(3, 27) = 6.09, p = 0.003), and alcohol intake (Table 2; F(3, 27) = 7.90, p < 0.001). Tukey’s post-hoc test revealed significantly reduced alcohol lever responses at the 5000 ng dose compared to vehicle (Figure 3B), and significantly reduced alcohol intake (g/kg) at both the 1000 and 5000 ng doses (Table 2). Analysis of cumulative alcohol lever responses across the session (Fig 3C) found significant main effects of drug (F(3, 27) = 5.37, p = 0.005), time (F(5, 45) = 59.5, p < 0.001), and a drug by time interaction (F(15, 135) = 5.40, p < 0.001). Post-hoc analysis showed that the 5000 ng dose significantly reduced responding relative to vehicle from minute 10 onwards and the 1000 ng dose significantly reduced responding from minute 15 onwards. There was no effect of eplerenone on inactive lever responses (Fig 3D) or locomotor rate (Fig 3E). Together these data suggest functional involvement of CeA MR, but not dHC MR in modulating alcohol self-administration.

**Figure 2.**
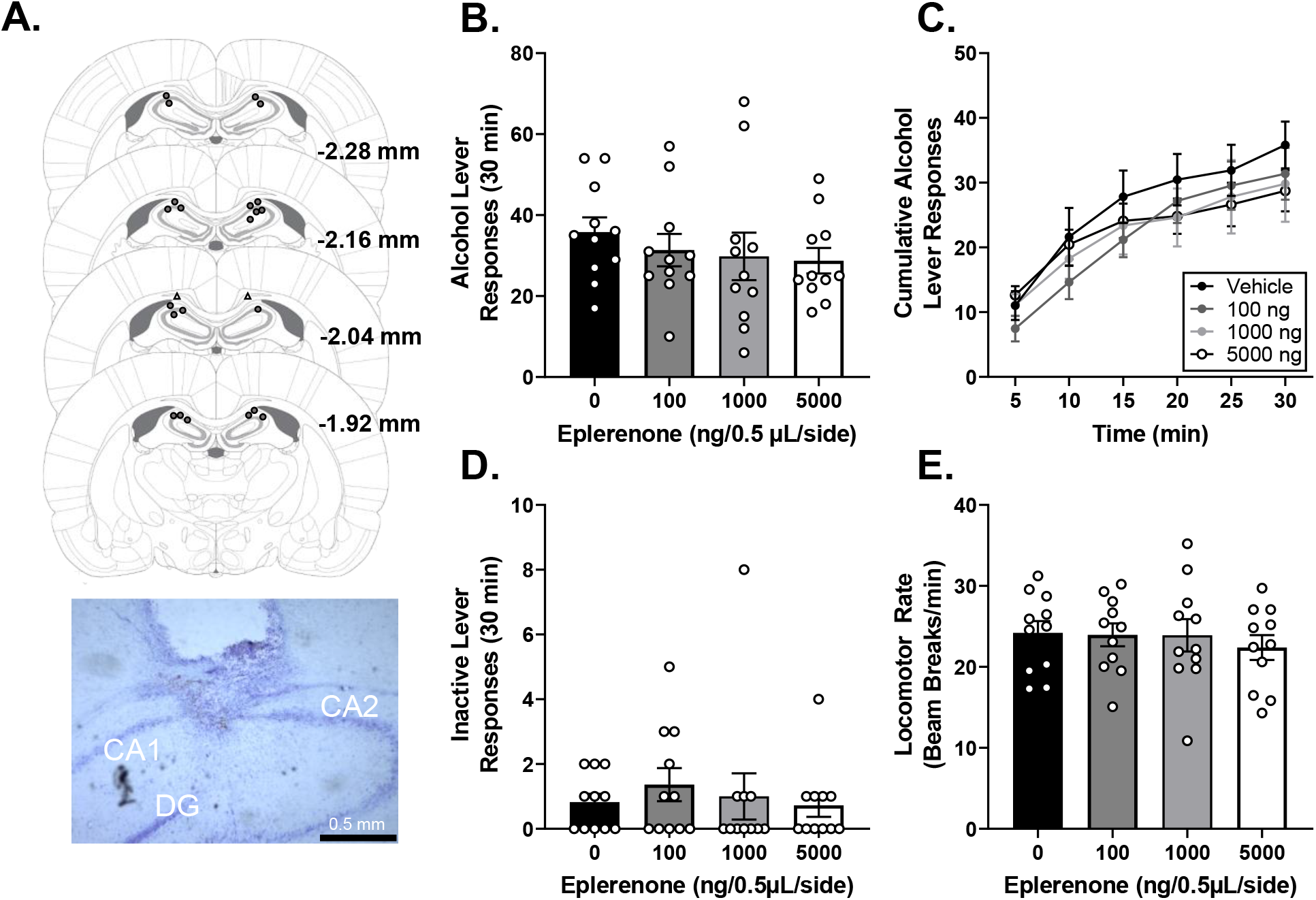
Intra-dHC eplerenone does not affect alcohol self-administration. (A) Bilateral dHC injector placements (hits represented by closed circles, misses represented by open triangles) and representative photomicrograph (4X). (B&C) Eplerenone does not reduce alcohol lever responses in female rats (n = 11). (D&E) Eplerenone has no effect on inactive lever responding or locomotor rate.

**Figure 3.**
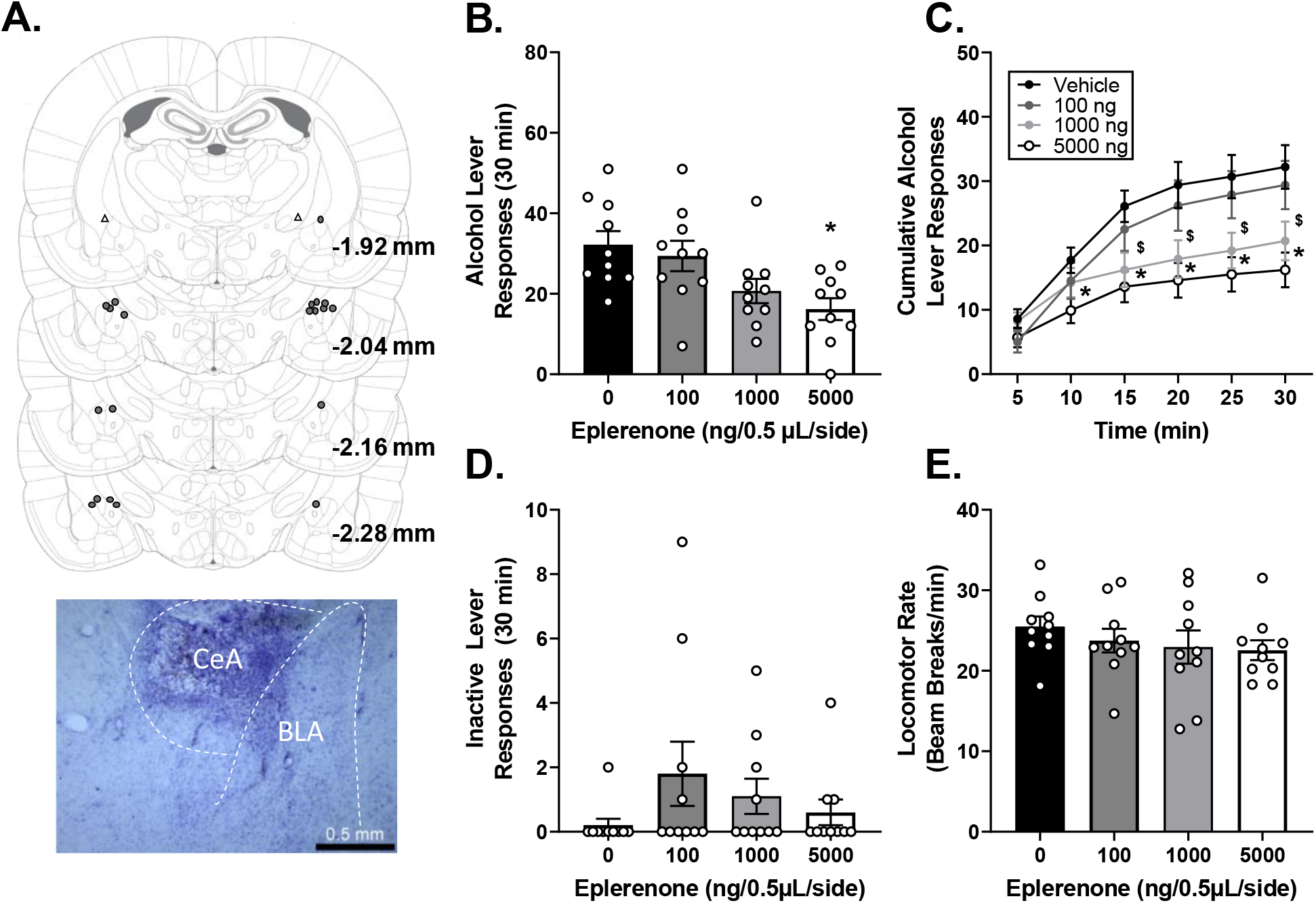
Intra-CeA eplerenone dose dependently reduces alcohol self-administration. (A) Bilateral CeA injector placements (hits represented by closed circles, misses represented by open triangles) and representative photomicrograph (4X). (B) Eplerenone (5000 ng) reduces alcohol lever responding in female rats (n = 10). (C) Eplerenone reduces alcohol lever responses from minute 10 onward (5000 ng) and 15 minutes onward (1000 ng). (D&E) Eplerenone has no effect on inactive lever responding or locomotor rate. *p < 0.05 eplerenone 5000 ng versus vehicle, ^$^p < 0.05 eplerenone 1000 ng versus vehicle.

**Table 2.** Alcohol intake for Experiment 2

| Eplerenone<br>(ng/0.5μL/side) | Alcohol Intake (g/kg) |  |  |  |
| --- | --- | --- | --- | --- |
|  | 0 | 100 | 1000 | 5000 |
| dHC | 0.81 ± 0.08 | 0.74 ± 0.09 | 0.68 ± 0.13 | 0.64 ± 0.08 |
| CeA | 0.78 ± 0.08 | 0.67 ± 0.07 | 0.47 ± 0.06* | 0.38 ± 0.06* |

\* - $p < 0.05$ versus 0 ng Eplerenone

### 3.3. Experiment 3: Confirmation of MR knockdown using ASO infusion

The purpose of this experiment was to confirm that ASO infusion would knock down MR expression, and to determine whether gene knockdown would persist for 7 days. Cannulae placement was visually verified prior to tissue punch and there were no misses.

#### 3.3.1. ASO infusion results in CeA MR knockdown 2 days post-infusion

CeA MR protein was quantified by western blot (Fig 4A). Welch’s ANOVA showed a main effect of ASO treatment (Fig 4B; W(2, 4.16) = 74.88, p < 0.001) on CeA MR protein expression, with the ASO-2D group showing significantly reduced CeA MR protein relative to the SCR group.

**Figure 4.**
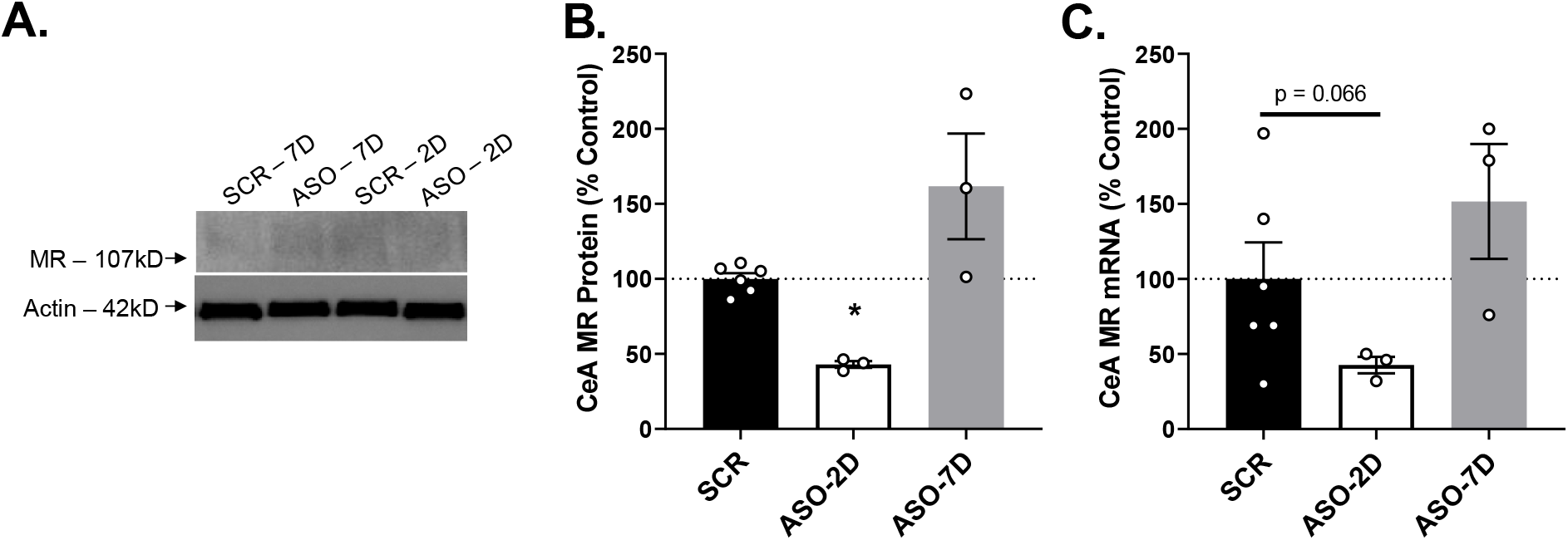
Verification of CeA MR Knockdown. (A) Representative western blot with bands for MR at 107kD and Actin at 42kD. (B) ASO treatment significantly reduced CeA MR protein expression 2 days post-infusion (n = 3/group ASO and 6/group SCR). (C) ASO treatment resulted in a trend for reduced CeA MR mRNA levels 2 days post-infusion. *p < 0.05 versus SCR.

There was a trend for a significant effect of ASO treatment on CeA MR mRNA (Fig 4C; p = 0.101). Given the western blot findings (Fig 4B), Welch’s t-test was used to compare CeA MR mRNA between the SCR and 2 day ASO groups and found a trend for decreased CeA MR mRNA between the groups (t(5.48) = 2.29, p = 0.066).

### 3.4. Experiment 4: Effect of central amygdala mineralocorticoid receptor knockdown on alcohol self-administration

The purpose of this experiment was to assess the functional role of CeA MR tone in alcohol self-administration.

#### 3.4.1. Knockdown of CeA MR results in a transient reduction of alcohol self-administration

Cannulae placement is shown in Fig 6B, 7 rats (3 in the SCR group and 4 in the ASO group) had one or both cannula(e) outside the target and were considered misses and excluded from analyses. Following ASO infusion Female rats showed a main effect of session and an ASO by session interaction on alcohol lever responses (Fig 6C; Session: F(4, 60) = 2.97, p = 0.027; Interaction: F(4, 60) = 2.76, p =0.036), with no main effect of ASO. Post-hoc analyses showed reduced alcohol lever responses in the ASO group relative to the SCR group 1 day post-infusion (p < 0.05). There was a main effect of session and a trend for ASO by session interaction on alcohol intake (Table 3; Session: F(4, 60) = 3.19, p = 0.019; Interaction: F(4, 60) = 2.28, p =0.071) indicating reduced alcohol intake in later sessions. There was no main effect of ASO. Post-hoc analysis showed lower alcohol intake in the ASO group relative to SCR on day 1 post-infusion (p < 0.05). There were no significant effects of ASO, session, or ASO by session interaction on inactive lever responses (Fig 6D). Locomotor rate showed a main effect of session (Fig 6E; F(4, 60) = 2.66, p = 0.041) with no effects of ASO or ASO by session interaction. These findings support those from Experiment 2 and suggest that CeA MR modulates alcohol self-administration in female rats.

**Table 3.**
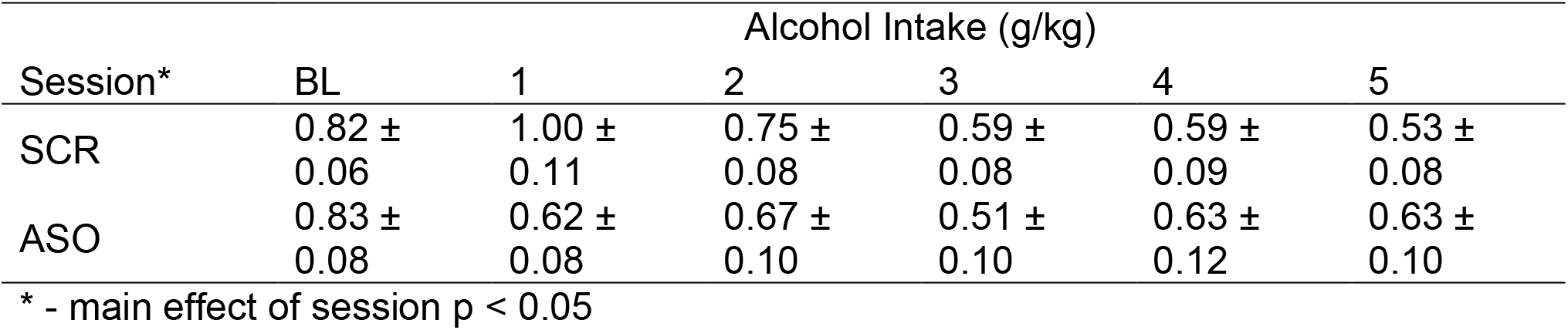
Alcohol intake for Experiment 3

## 4. DISCUSSION

The results of this study provide further evidence for the functional role of MR in modulating alcohol consumption. First, it was confirmed that systemic MR antagonism with spironolactone increases plasma corticosterone while reducing alcohol self-administration in female rats, and that the reductions in self-administration correlate with increased plasma corticosterone. Next, local microinjections demonstrated that CeA but not dHC MR antagonism reduced alcohol self-administration in female rats. Finally, it was shown that ASO knockdown of CeA MR transiently reduced alcohol self-administration in female rats. Together, these data suggest that CeA MRs modulate alcohol self-administration and could pose an interesting target for pharmacological treatment of AUD.

As MR regulate basal levels of the hormone corticosterone (Joels and de Kloet, 2017) which is known to modulate alcohol self-administration (Fahlke et al., 1994a, Fahlke et al., 1994b), we hypothesized that the observed reduction in alcohol self-administration following spironolactone pretreatment (Makhijani et al., 2018) was due to stimulation of the HPA axis by inhibition of glucocorticoid negative feedback (Atkinson et al., 2008). Indeed, systemic administration of spironolactone (50 mg/kg) increased plasma corticosterone similar to other MR antagonists (Ratka et al., 1989, Bitran et al., 1998). Furthermore, there was significant correlation between reduced self-administration and increased corticosterone, suggesting a relationship between these two effects. This relationship is similar to the purported mechanism of action for naltrexone (O’Malley et al., 2002, Kiefer et al., 2006), where stimulation of the HPA axis reduces alcohol craving, and is also the mechanism by which MR antagonists mediate anxiolytic effects (Bitran et al., 1998). In contrast, Experiment 2 also showed reduced alcohol self-administration following intra-CeA infusion of the MR antagonist eplerenone, which is not known to impact corticosterone levels when injected systemically (Hlavacova and Jezova, 2008, Hlavacova et al., 2010). Therefore, it is possible that increased corticosterone is immaterial to the observed reductions in self-administration; however, as we did not assess corticosterone levels following eplerenone infusion in this study we cannot definitively exclude the role of increased corticosterone. An alternative explanation is that the effects of MR antagonism on alcohol self-administration are mediated by different peripheral versus central mechanisms (i.e. corticosterone may mediate the effects of peripheral MR antagonism but not intra-CeA antagonism). While Experiment 1 adds to our previous findings in elucidating a potential mechanism of action for the systemically administered MR antagonist modulation of alcohol self-administration, there remain several caveats such as the contribution of peripheral versus central MR effects (Jaisser and Farman, 2016), and off target spironolactone effects (Makhijani et al., 2018).

To address these questions, Experiment 2 utilized regional microinjections of the more selective MR antagonist eplerenone (de Gasparo et al., 1987). Infusion of eplerenone into the CeA dose-dependently decreased alcohol self-administration while dHC infusion had no effect on alcohol self-administration. This finding is in agreement with literature suggesting CeA MR are involved in alcohol drinking (Aoun et al., 2018), and dHC MR are involved in other behaviors including response to novelty, spatial memory, and anxiety-like behavior (Oitzl and de Kloet, 1992, Lai et al., 2007, McCann et al., 2019). Interestingly, while MR is implicated in regulating memory (Ninomiya et al., 2010, Zhou et al., 2010, McCann et al., 2019) the reduction in alcohol self-administration in the present experiments is likely not due to a memory impairment as there was no alteration in discrimination between the alcohol and inactive levers (i.e., greater responses on the inactive lever relative to the alcohol lever). In addition to identifying the brain locus of MR effects on drinking, this study increases confidence that MR mediates these effects on alcohol self-administration by using the more selective MR antagonist eplerenone instead of spironolactone (de Gasparo et al., 1987). Furthermore, this study increases the translational relevance of these findings as eplerenone has a greater safety profile than spironolactone, which has an FDA black box warning (Lainscak et al., 2015).

To directly assess the impact of CeA MR tone on alcohol self-administration, a validated ASO construct (Sakai et al., 2000, Johnson and Greenwood-Van Meerveld, 2015) was infused into the CeA of female rats prior to alcohol self-administration. Consistent with our studies utilizing an MR antagonist (Experiments 2 & 3; 3.1. & 3.2.), rats showed a transient decrease in alcohol self-administration 1 day after ASO infusion. The transient nature of the reduction coupled with confirmed MR knockdown 2 days post-infusion (Fig 4) suggests a compensatory mechanism restoring normal self-administration behavior. One possible explanation is that reduced self-administration is due to a transient increase in corticosterone following MR knockdown, as in Experiment 1 (3.1.2.). This is supported by the fact that corticosterone is increased acutely following MR antagonism (Fig 1), but not in genomic models of altered MR expression, indicating the presence of compensatory mechanisms for regulating corticosterone in the absence of normal MR function (Rozeboom et al., 2007). It is also possible that this transient reduction is due to off-target effects of the ASO infusion and not MR knockdown, as we did not assess the level of MR knockdown 1 day after ASO infusion, when the behavioral effect was observed. However, this limitation is tempered by previous studies reporting biological effects of MR ASO knockdown 1 day post-infusion (Ma et al., 1997, Sakai et al., 2000), and the presence of a scrambled oligonucleotide control. Future studies could expand upon this hypothesis by examining corticosterone levels at multiple timepoints following CeA MR knockdown.

Interestingly, the directionality of both MR antagonism and pharmacological knockdown effects on alcohol self-administration differ from those described in Aoun et al., 2018. While Aoun et al. showed lower levels of MR expression and higher levels of aldosterone correlate with higher drinking behaviors across multiple species, here we demonstrate that antagonism and pharmacological knockdown of CeA MR reduces alcohol self-administration. This paradoxical disconnect between MR tone and pharmacological manipulation is not unique to these self-administration studies, as there is a similar inverse relationship between MR expression and anxiety-like/fear behavior. For example, genomic MR knockdown reduces anxiety-like/fear behavior (Brinks et al., 2009, McCann et al., 2019) and genomic overexpression ameliorates these behaviors (Lai et al., 2007, Rozeboom et al., 2007, Ter Horst et al., 2012); however, MR antagonists are shown to have anxiolytic effects (Bitran et al., 1998, Hlavacova and Jezova, 2008, Hlavacova et al., 2010, Lopez-Rubalcava et al., 2013). Possible explanations for this phenomenon include differences in intra-CeA distribution of MR expression and genomic versus non-genomic effects of MR. While the MR is known to be expressed throughout limbic brain systems (McEwen et al., 1968, Reul and de Kloet, 1985), little is known about its expression pattern within the CeA, a region with complex microcircuitry (Gilpin et al., 2015). Differences in MR expression that impact alcohol drinking may be biased in localization towards GABAergic interneurons, projection neurons, or glutamatergic afferents, while the effect of an antagonist or ASO would presumably be uniform which may explain these conflicting findings. Alternatively, while the timing of MR antagonist effects on alcohol self-administration (within minutes) suggest that they are non-genomically mediated (Karst et al., 2005, Khaksari et al., 2007), the effect of MR tone on alcohol drinking may be mediated, partially or in full, by genomic MR action as a ligand-dependent transcription factor (Fuller et al., 2000, Ruhs et al., 2017). Notably, the ASO knockdown utilized in these studies was transient and resulted in a similar behavioral effect as MR antagonism on self-administration, while knockdown studies finding opposite results to antagonism utilize genomic manipulation of MR (Cobden et al., 1988, Lai et al., 2007, Rozeboom et al., 2007, Brinks et al., 2009, Ter Horst et al., 2012). Differences in the impact of MR expression and antagonism on alcohol self-administration could be further clarified by characterizing MR expression patterns within the CeA of animals with high and low MR expression, or by assessing alcohol self-administration in animals with genomically altered CeA MR expression.

Together, these results confirm our previous findings that systemic MR antagonism reduces alcohol self-administration, proposes a potential mechanism of action involving increased plasma corticosterone, and demonstrates the functional role of CeA MR in alcohol self-administration utilizing the selective MR antagonist eplerenone and ASO knockdown. The effects of MR antagonism on alcohol self-administration are particularly relevant as eplerenone is also currently used for treatment of hypertension and cardiovascular disease (Stewart Coats and Shewan, 2015) which are exacerbated by and comorbid with AUD (Gardner and Mouton, 2015). Additionally, as pharmacological management of these conditions is well accepted, use of MR antagonists to treat comorbid cardiovascular disease and AUD may be a novel approach to overcoming the AUD treatment gap (Poulter et al., 2015, Whelton et al., 2018). Future studies could further explore the clinical implications of this data by analyzing alcohol drinking patterns in patients prescribed eplerenone, and expand upon the molecular and circuit implications by clarifying the role of genomic versus nongenomic MR signaling in alcohol self-administration, or by characterizing the localization of MR within CeA circuitry. Altogether, this study adds to the growing body of literature suggesting that MR plays a role in corticosteroid regulation of alcohol drinking, and presents a potential new target for pharmacological treatment of alcohol use disorder.

**Figure 5.**
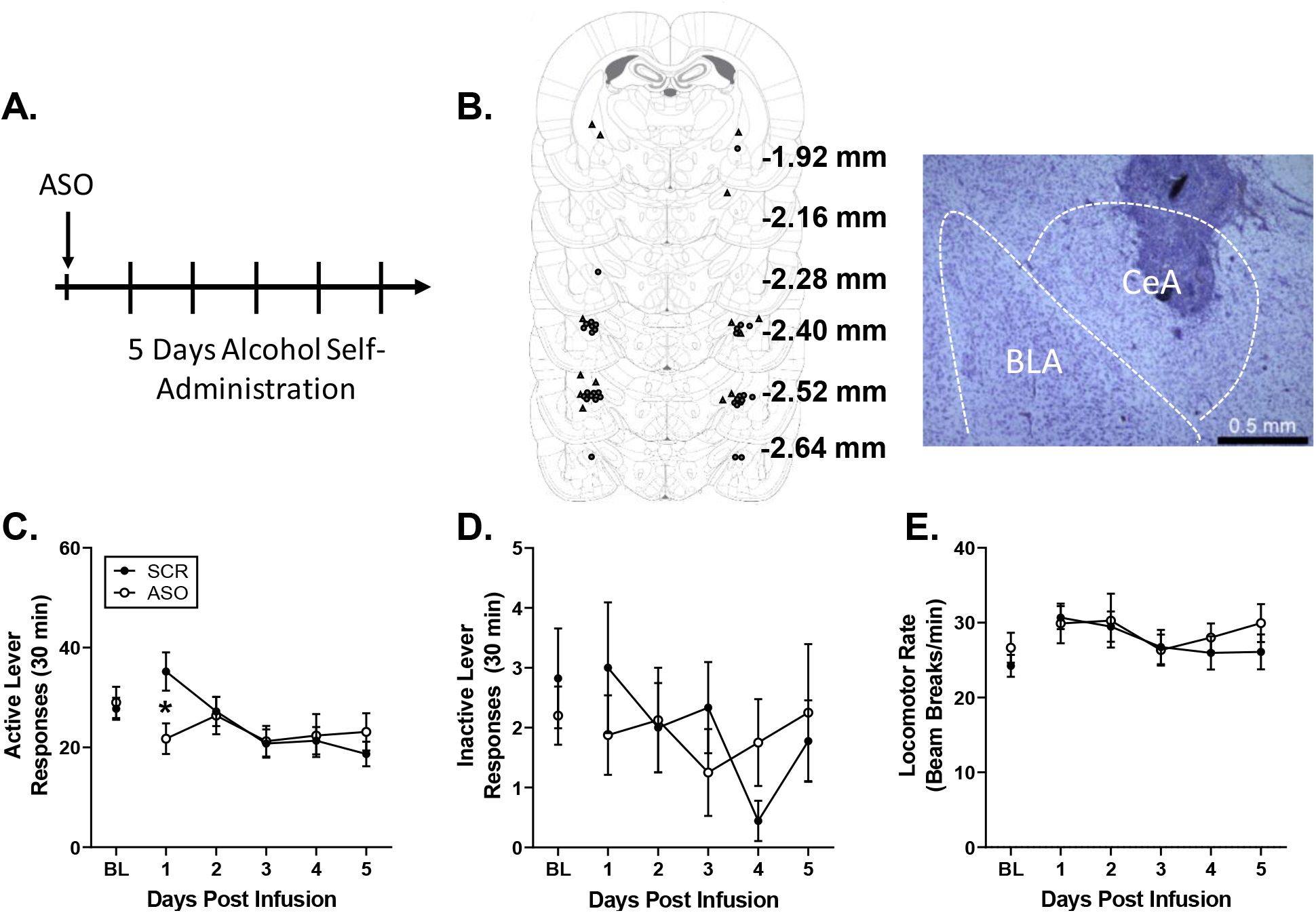
Transient reduction in alcohol self-administration following CeA MR knockdown in female rats. (A) Timeline depicting dates of ASO infusion and alcohol self-administration testing. (B) Bilateral CeA cannulae placements (hits represented by closed circles, misses represented by open triangles) and representative photomicrograph (4X). (C) CeA MR knockdown results in a transient reduction in alcohol self-administration in female rats (n = 9/SCR group and 8/ASO group). (D-E) CeA MR knockdown had no effect on inactive lever responses, or locomotor rate. *p < 0.05 versus SCR.

## Acknowledgements

This work was supported in part by the National Institute of Health AA026537 (JB) and by the Bowles Center for Alcohol Studies. VHM was supported by AA027436 and NS007431.

